## Supplementary material for "Structural insights into p300 regulation and acetylation-dependent genome organisation": histones MS

**Protein Modification Analysis**

Table of Contents

Protein 1, H4	2

```
Protein 2, H2A   	3
Protein 3, H2B  	3
```

Protein 4, H3	4

Matched peptides shown in **Bold Red**

**His6-H4**

1 MKSSHHHHHH ENLYFQSNAM SGR**GKGGKGL GKGGAKR**HRK **ILRDNIQGIT KPAIRR**LARR

61 GGVK**RISGLI YEEVRAVLKS FLESVIRDSV TYTEHAKRKT VTSLDVVYAL KR**QGR**TLYGF GG**

Sequence coverage (regardless of tag): 83%

**Quantification results for acetylation (K)**

S1: Sample 1

S2: Sample 2

“Intensity”: indicates the normalized signal for each peptide of interest.

| **PTM** | **Peptide Sequence** | **Intensity CoreNCP, S1** | **Intensity**  **CoreNCP, S2** | **Intensity**  **Only NCP, S1** | **Intensity**  **Only NCP, S2** | **Intensity**  **P300sNCP, S1** | **Intensity**  **P300sNCP, S2** |
| --- | --- | --- | --- | --- | --- | --- | --- |
| 1xAcetyl [K12] | [K].TVTSLDVVYALKR.[Q] | 144.6 | 172.3 | NA | NA | 164.1 | 119 |
| 1xAcetyl [K1] | [R].KTVTSLDVVYALK.[R] | 144.4 | 150.6 | NA | NA | 156.6 | 148.4 |
| 1xAcetyl [K10] | [R].DSVTYTEHAKR.[K] | 137.4 | 134.8 | NA | NA | 164.3 | 163.6 |
| 1xAcetyl [K8] | [R].DNIQGITKPAIR.[R] | 155.6 | 137.4 | 19.1 | NA | 127.4 | 160.5 |
| 2xAcetyl [K3; K7] | [K].GGKGLGKGGAK.[R] | 268.9 | 312.7 | NA | NA | 12.8 | 5.6 |
| 2xAcetyl [K4; K8] | [K].GLGKGGAKR.[H] | 251.1 | 340.9 | NA | NA | 8 | NA |
| 4xAcetyl [K2; K5; K9; K13] | [R].GKGGKGLGKGGAKR.[H] | 113.6 | 159 | 16.7 | NA | 191.4 | 119.4 |
| 3xAcetyl [K5; K9; K] | [R].GKGGKGLGKGGAKR.[H] | 234.1 | 365.9 | NA | NA | NA | NA |
| 3xAcetyl [K2; K5; K9] | [R].GKGGKGLGKGGAK.[R] | 210.4 | 296.7 | 9 | NA | 51.3 | 32.5 |
| 2xAcetyl [K5; K9] | [R].GKGGKGLGKGGAK.[R] | 248.1 | 311.3 | NA | 22.2 | 18.4 | NA |
| 2xAcetyl [K2; K5] | [R].GKGGKGLGK.[G] | 258.6 | 287.2 | 4.2 | 9.7 | 29.7 | 10.7 |
| 1xAcetyl [K5] | [R].GKGGKGLGK.[G] | 80.5 | 341.4 | 6.8 | 19.9 | 104.5 | 46.8 |
| 3xAcetyl [K3; K7; K11] | [K].GGKGLGKGGAKR.[H] | 273.8 | 326.2 | NA | NA | NA | NA |
| 2xAcetyl [K3; K7] | [K].GGKGLGKGGAKR.[H] | 287.4 | 312.6 | NA | NA | NA | NA |

**His6-H2A**

1 MKSSHHHHHH ENLYFQSNAM SGMSGGK**GGK AGSAAKASQS RSAKAGLTFP VGR**VHRLLR**R**

**61 GNYAQRIGSG APVYLTAVLE YLAAEILELA GNAAR**DNKKT RIIPR**HLQLA IRNDDELNKL**

**121 LGNVTIAQGG VLPNIHQNLL PK**KSAKATKA SQEL

Sequence coverage (regardless of tag): 74%

**Quantification results for acetylation (K)**

S1: Sample 1

S2: Sample 2

“Intensity”: indicates the normalized signal for each peptide of interest.

| **PTM** | **Peptide Sequence** | **Intensity CoreNCP, S1** | **Intensity**  **CoreNCP, S2** | **Intensity**  **Only NCP, S1** | **Intensity**  **Only NCP, S2** | **Intensity**  **P300sNCP, S1** | **Intensity**  **P300sNCP, S2** |
| --- | --- | --- | --- | --- | --- | --- | --- |
| 2xAcetyl [K3; K9] | [K].GGKAGSAAKASQSR.[S] | NA | NA | NA | NA | NA | NA |
| 1xAcetyl [K3] | [R].SAKAGLTFPVGR.[V] | 135.1 | 217 | NA | NA | 135.3 | 112.6 |
| 1xAcetyl [K6] | [K].AGSAAKASQSR.[S] | 185.2 | 154.8 | NA | NA | 179.7 | 80.2 |

**H2B**

The peptide sequence received is shown as below, which was not detected in the sample.

1 MNFSSRDSRL LVIRSQFRSF SSDSFKDIV NERVQDRHGL LGNTSVRVSL LQNFVNVRRV

61 CFLSSLASLL TISGSGSLFS GWFLFSWGF GGWFLFGFSR HMYISFLKLN KIISRASEFQ

121 LELSGCEIVI RWYQA

However, yeast histone H2B.1 was detected with high coverage. The sequence is shown as below:

**1 MSAKAEKKPA SKAPAEKKPA AKKTSTSTDG KKR**SKAR**KET YSSYIYKVLK QTHPDTGISQ**

**61 KSMSILNSFV NDIFERIATE ASKLAAYNKK STISAREIQT AVRLILPGEL AKHAVSEGTR**

**121 AVTK**YSSSTQ A

Sequence coverage (regardless of tag): 92%

**Quantification results for acetylation (K)**

S1: Sample 1

S2: Sample 2

“Intensity”: indicates the normalized signal for each peptide of interest.

| **PTM** | **Peptide Sequence** | **Intensity CoreNCP, S1** | **Intensity**  **CoreNCP, S2** | **Intensity**  **Only NCP, S1** | **Intensity**  **Only NCP, S2** | **Intensity**  **P300sNCP, S1** | **Intensity**  **P300sNCP, S2** |
| --- | --- | --- | --- | --- | --- | --- | --- |
| 1xAcetyl [K9] | [R].LILPGELAKHAVSEGTR.[A] | 205.3 | 130.2 | 68.4 | NA | 91.3 | 104.8 |
| 1xAcetyl [K7] | [R].IATEASKLAAYNK.[K] | 197.1 | 144.9 | NA | NA | 91.2 | 166.8 |
| 1xAcetyl [K3] | [K].VLKQTHPDTGISQK.[S] | 182.3 | NA | 18.9 | NA | 224.8 | 173.9 |
| 2xAcetyl [K4; K7]; 1xMet-loss [N-Term] | [-].MSAKAEKKPASK.[A] | 144.6 | 182.1 | NA | NA | 173.8 | 99.5 |
| 2xAcetyl [K1; K9] | [K].KTSTSTDGKKR.[S] | 600 | NA | NA | NA | NA | NA |
| 3xAcetyl [K6; K10; K11] | [K].APAEKKPAAKKTSTSTDGK.[K] | 151.6 | 153.7 | 38.3 | NA | 256.4 | NA |
| 2xAcetyl [K6; K10] | [K].APAEKKPAAKK.[T] | 157.4 | 278.8 | NA | NA | 88.3 | 75.5 |
| 1xAcetyl [K] | [K].APAEKKPAAK.[K] | 239 | 229.7 | 3.9 | NA | 84.1 | 43.2 |
| 3xAcetyl [K4; K8; K13] | [K].AEKKPASKAPAEKKPAAK.[K] | 150 | 391.4 | NA | NA | 58.6 | NA |
| 2xAcetyl [K4; K8] | [K].AEKKPASKAPAEK.[K] | 136.6 | 246.5 | 19 | NA | 107.4 | 90.5 |
| 1xAcetyl [K8] | [K].AEKKPASKAPAEK.[K] | 191.4 | 382.3 | NA | NA | NA | 26.2 |
| 2xAcetyl [K5; K10] | [K].KPASKAPAEKKPAAK.[K] | 109.8 | 220.2 | 2.6 | NA | 163.4 | 103.9 |
| 1xAcetyl [K5] | [K].KPASKAPAEK.[K] | 241.4 | 218.1 | 5.5 | NA | 80.4 | 54.6 |
| 2xAcetyl [K5; K6] | [K].KPAAKKTSTSTDGK.[K] | 132.6 | 129.8 | NA | NA | 204.5 | 133.1 |
| 1xAcetyl [K1] | [R].KETYSSYIYK.[V] | 150.6 | 117.7 | 34.3 | NA | 146 | 151.4 |

**His-H3**

1 MKSSHHHHHH ENLYFQSNAM ARTKQTAR**KS TGGKAPRKQL ASKAARKSAP STGGVKKPHR**

**61 YKPGTVALR**E IRR**FQKSTEL LIRKLPFQR**L VR**EIAQDFKT DLRFQSSAIG ALQESVEAYL**

**121 VSLFEDTNLA AIHAK**RVTIQ KKDIKLARRL RGERS

Sequence coverage (regardless of tag): 74%

**Quantification results for acetylation (K)**

S1: Sample 1

S2: Sample 2

“Intensity”: indicates the normalized signal for each peptide of interest.

| **PTM** | **Peptide Sequence** | **Intensity CoreNCP, S1** | **Intensity**  **CoreNCP, S2** | **Intensity**  **Only NCP, S1** | **Intensity**  **Only NCP, S2** | **Intensity**  **P300sNCP, S1** | **Intensity**  **P300sNCP, S2** |
| --- | --- | --- | --- | --- | --- | --- | --- |
| 1xAcetyl [K5] | [K].STGGKAPR.[K] | 180.8 | 366.8 | NA | NA | 36.8 | 15.6 |
| 1xAcetyl [K9] | [K].SAPSTGGVKKPHR.[Y] | 116 | 198.4 | NA | NA | 91.7 | 193.9 |
| 1xAcetyl [K5] | [K].QLASKAAR.[K] | 295.1 | 250.7 | NA | NA | 33.6 | 20.6 |
| 1xAcetyl [K1] | [R].KSAPSTGGVK.[K] | 258.7 | 253.3 | 4.6 | 5.8 | 48.9 | 28.5 |
| 1xAcetyl [K] | [R].KSTGGKAPR.[K] | 20.8 | 226.5 | 65.9 | 233.1 | 13.8 | 39.9 |
| 1xAcetyl [K7] | [R].EIAQDFKTDLR.[F] | 172.1 | 161.4 | 43.4 | NA | 98.9 | 124.2 |
| 1xAcetyl [K3] | [R].FQKSTELLIR.[K] | 120.7 | 105.6 | 169.1 | NA | 91.1 | 113.5 |
| 2xAcetyl [K1; K10] | [R].KSAPSTGGVKKPHR.[Y] | 218.4 | 256.1 | NA | NA | 125.4 | NA |
| 2xAcetyl [K1; K6] | [R].KQLASKAAR.[K] | 248.2 | 317.8 | 2.8 | 3.8 | 27.3 | NA |
| 1xAcetyl [K] | [R].KQLASKAAR.[K] | 221.9 | 242.5 | 7.1 | 18.7 | 53.8 | 56 |
| 2xAcetyl [K1; K6] | [R].KSTGGKAPR.[K] | 232.8 | 296.5 | 2.9 | 5.5 | 34.7 | 27.6 |
| 1xAcetyl [K2] | [R].YKPGTVALR.[E] | 138.7 | 134.5 | NA | NA | 151.5 | 175.3 |
